# Engineering *Clostridium thermocellum* for production of 2,3-butanediol from cellulose

**DOI:** 10.1101/2025.11.28.691234

**Authors:** S. Bilal Jilani, Nandhini Ashok, Yannick J. Bomble, Adam M. Guss, Daniel G. Olson

## Abstract

*Clostridium thermocellum* is a promising host for consolidated bioprocessing due to its ability to directly ferment cellulose into fuels and chemicals. However, natural product formation in this organism is limited. Here, we report engineering *C. thermocellum* for the production of 2,3-butanediol (23BD), a valuable industrial chemical. We functionally expressed a thermophilic 23BD pathway in this organism resulting in a 23BD titer of 19.7 mM from cellulose, representing a metabolic yield of 24%. We used a cell-free systems biology approach to identify limiting steps in the 23BD pathway, revealing that exogenous 23BD dehydrogenase (BDH) activity was essential for production, while native acetolactate synthase (ALS) and acetolactate decarboxylase (ALDC) activities were present but limiting in the parent strain. This approach also revealed redox balance limitations. We demonstrated that this improved understanding of redox balance limitations could be used to increase 23BD titer in vivo, showing that adding acetate could be used to increase 23BD yield. This work establishes a foundation for developing *C. thermocellum* into a robust platform for 23BD production directly from cellulose and highlights the utility of cell-free systems for guiding metabolic engineering in non-model organisms.

## Introduction

Cellulosic biomass is an abundant and renewable feedstock that could be used to produce a variety of fuels and chemicals; despite this, its recalcitrance to microbial conversion has limited commercial applications so far (Lynd, 2017; Lynd et al., 2022). Native cellulolytic organisms provide one possible solution to the problem of recalcitrance; yet, these organisms do not naturally produce fuels and chemicals at high yield or titer. One native cellulolytic organism that has been the focus of substantial prior work is *Clostridium thermocellum* (recently renamed *Hungateiclostridium thermocellum*, *Ruminiclostridium thermocellum*, and now *Acetivibrio thermocellus* (Tindall, 2019)). It is able to achieve rapid solubilization of lignocellulosic biomass without the need for thermochemical pretreatment (Holwerda et al., 2019, 2013) and thus reduces the formation of microbial metabolic inhibitors from the chemical/thermal breakdown of biomass components (Jilani and Olson, 2023). This rapid solubilization is achieved due to a specialized multi-enzyme protein complex known as a cellulosome (Bayer et al., 2008). To take advantage of its native cellulolytic capabilities, product formation needs to be increased. Much prior work in *C. thermocellum* has focused on increasing ethanol production (Mazzoli and Olson, 2020); however, product titer is currently limited by the toxicity of ethanol (Tian et al., 2017). Since *C. thermocellum* is a promising microbial chassis for CBP, it could potentially be used as a biocatalyst for production of other commodity chemicals.

2,3-butanediol (23BD) is an industrial chemical that has been used as a plasticizer, pharmaceutical ingredient, agricultural chemical, and precursor for synthetic rubber (Lee et al., 2019; Straathof, 2014). It has two chiral centers, and three separate stereoisomers can be produced: (S, S)-, (R, R)-, and meso-23BD. Among the three, the (R, R) form is the most commercially valuable, and a further benefit of biological production is the ability to produce chirally pure forms (Ji et al., 2015). It is an aliphatic polyol, a category of chemicals that are notable for their low microbial toxicity, enabling production of polyols at very high titers: 300 g/L glycerol (Vijaikishore and Karanth, 1987), 131 g/L for 1,3-propanediol (Cervin et al., 2010), 125 g/L 1,4-butanediol (Burgard et al., 2016), and 178 g/L for 23BD (Lee and Seo, 2019).

There are two primary challenges associated with 23BD production in a thermophilic anaerobic organism such as *C. thermocellum*. The first is that there are few well characterized thermophilic candidate enzymes for several reactions needed to convert pyruvate to 23BD. The most well-characterized pathway consists of three enzymes: acetolactate synthase (ALS), acetolactate decarboxylase (ALDC), and 23BD dehydrogenase (BDH), and several mesophilic isoenzymes for each reaction have been identified (Kay and Jewett, 2015; Ma et al., 2009; Zhang et al., 2014). The moderately thermophilic *Bacillus licheniformis* has been shown to produce 23BD up to 55°C, suggesting that the corresponding genes may function in *C. thermocellum*. Two BDH enzymes have been identified in *B. licheniformis*: one that produces R,R-23BD (known as Gdh) and one that produces meso-23BD (known as BudA or BudC) (Ge et al., 2016). The genome of *B. licheniformis* is also predicted to encode a fermentative ALS enzyme (AlsS) and an ALDC enzyme (AlsD).

In *C. thermocellum*, the ALS reaction is thought to be present due to high levels of valine production in some strains (Rydzak et al., 2017; van der Veen et al., 2013), and two genes (Clo1313_0099 and Clo1313_0305) are annotated as acetolactate synthase genes. Furthermore, a moderately thermostable AlsS was identified in *Bacillus subtilis* that functions in *C. thermocellum* at temperatures up to 50 °C (Lin et al., 2015, 2014). Thermophilic versions of the ALDC and BDH enzymes (Peretz et al., 1997; Xiao et al., 2012; Yu et al., 2011; Zheng et al., 2022) have not yet been expressed in *C. thermocellum*; however, low levels of 23BD (0.42 g/L) have been observed in *C. thermocellum* grown in the presence of very high substrate concentrations (97 g/L Avicel microcrystalline cellulose) (Holwerda et al., 2014), suggesting that low levels of these activities may be present in this organism. No putative acetolactate decarboxylase genes have been identified in *C. thermocellum*. Identifying *bdh* genes can be more challenging, since many organisms contain a variety of poorly annotated dehydrogenase genes. *C. thermocellum* has at least three possible candidate genes annotated as short-chain dehydrogenase/reductases: Clo1313_0815 (which has no additional genomic context), Clo1313_1283 (located in a fatty acid biosynthesis gene cluster), and Clo1313_2272 (which also has no additional genomic context).

A second challenge associated with 23BD production in *C. thermocellum* is that conversion of six-carbon (C6) sugars or sugar-equivalents to 23BD is not redox balanced. In glycolysis, conversion of one C6 sugar (or equivalent) to two three-carbon (C3) pyruvate molecules results in production of two reducing equivalents (i.e., NAD(P)H), only one of which is reoxidized during 23BD biosynthesis (Figure 1). In aerobic organisms, this redox imbalance can be addressed by careful control of aeration during growth (Boecker et al., 2021; Lee and Seo, 2019; Wichmann et al., 2023). Excess NADH can either be reduced with a water-forming NADH oxidase or used to generate ATP via the electron transport chain. However, both of these options require oxygen and are not possible in an obligate anaerobe such as *C. thermocellum*.

**Figure 1.**
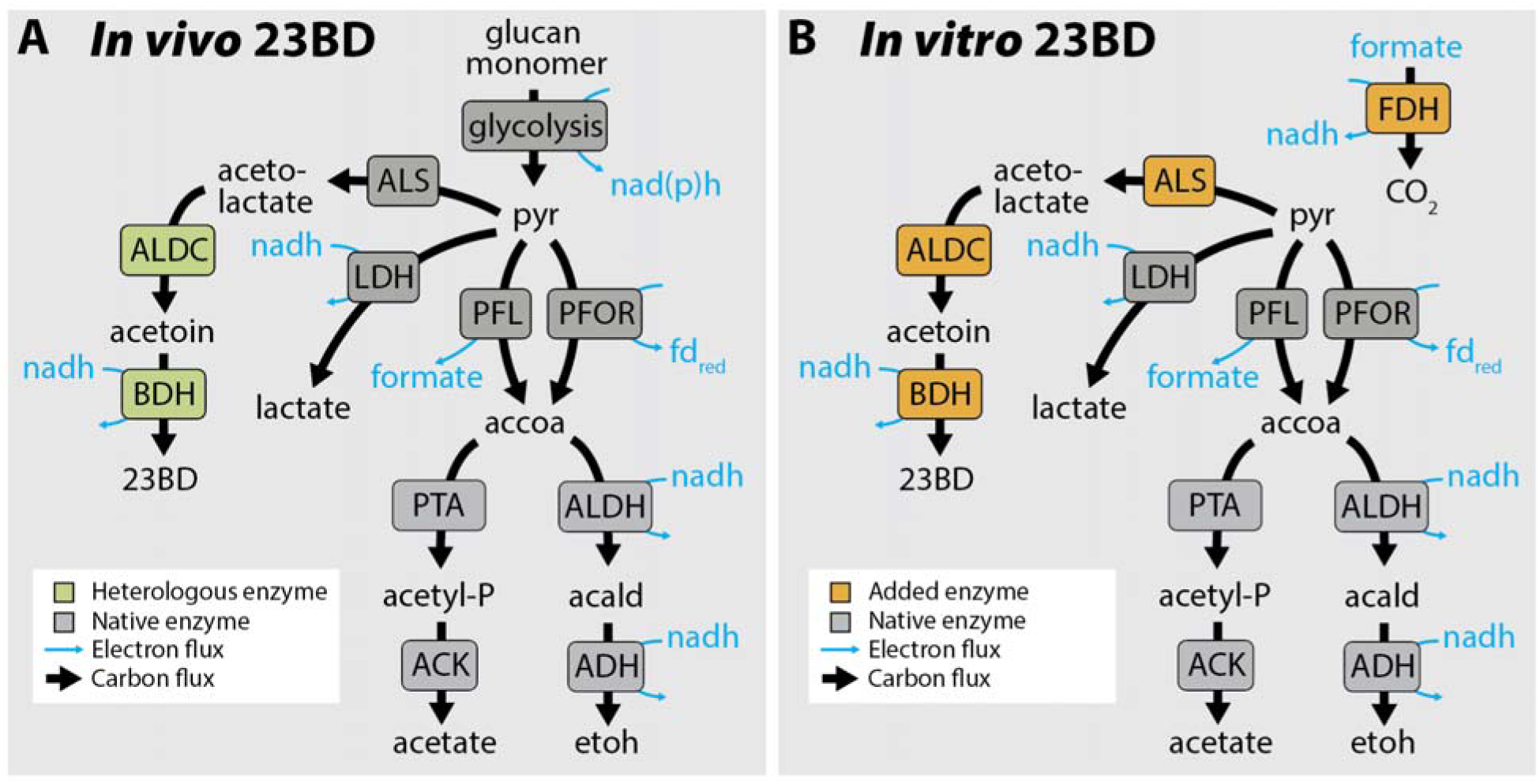
Pathways for 23BD production. **A.** Metabolic pathways for conversion of cellulose and soluble sugars to 23BD and other fermentation products in vivo. Heterologous enzymes are shown in green. Native enzymes are shown in grey. **B.** Metabolic pathways for conversion of pyruvate to 23BD and other fermentation products in the cell-free system. Enzymes added *in vitro* are shown in orange. Native enzymes are shown in grey. For both panels, black arrows indicate carbon fluxes, and blue arrows indicate electron fluxes. Enzyme abbreviations include: acetolactate synthase (ALS), acetolactate decarboxylase (ALDC), 2,3-butanediol dehydrogenase (BDH), lactate dehydrogenase (LDH), pyruvate formate lyase (PFL), pyruvate-ferredoxin oxidoreductase (PFOR), phosphotransacetylase (PTA), acetate kinase (ACK), acetaldehyde dehydrogenase (ALDH), alcohol dehydrogenase (ADH), formate dehydrogenase (FDH). Metabolite abbreviations include: pyruvate (pyr), 2,3-butanediol (23BD), acetyl-CoA (accoa), acetyl-phosphate (acetyl-P), acetaldehyde (acald), ethanol (etoh), reduced ferredoxin (fd_red_). For simplicity, only reduced cofactors are shown. The cofactor specificity of glycolysis is shown as nad(p)h to indicate that both NADH and NADPH can be produced by *C. thermocellum* glycolysis, depending on the relative flux between the pyruvate phosphate dikinase and malate shunt reactions.

Thus, the overall goals of this work were to (1) identify a thermophilic pathway for 23BD production that can be functionally expressed in *C. thermocellum*, and (2) identify and overcome redox imbalances associated with 23BD production from C6 sugars. By understanding the current limitations of the *C. thermocellum* 23BD pathway, we lay the foundation for further engineering efforts to make 23BD at high titer, rate, and yield.

## Materials and Methods

### Strains, plasmids, and culturing conditions

#### Cultivation conditions

For genetic manipulations, *C. thermocellum* strains were grown on rich CTFUD medium (Olson and Lynd, 2012). Medium was supplemented with 15 µg/mL thiamphenicol (Tm15) when needed, to select for plasmids. For measurement of fermentation products, *C. thermocellum* strains were cultivated in MTC-5 chemically defined medium (Cui et al., 2020) with either 5 or 20 g/L cellobiose in either 15 or 50 mL tubes at 55 °C under anaerobic conditions. Growth of *C. thermocellum* was performed in anaerobic conditions using a Coy Labs anaerobic chamber. A gas mix of 85% N_2_, 10% CO_2_, and 5% H_2_ was used to maintain the anaerobic conditions inside the chamber.

#### Strains

The parent strain used in this work, *C. thermocellum* LL1299, is a modified version of WT *C. thermocellum* DSM 1313. It has 2 genetic alterations: a deleted *hpt* gene to allow counter selection with 8-azahypoxantine (Argyros et al., 2011; Olson and Lynd, 2012) and a deleted restriction enzyme (Clo1313_0478) to increase transformation efficiency (Tian et al., 2019). This strain was further modified to create strain AG8235 by insertion of a “landing pad” sequence by homologous recombination (Elmore et al., 2023) to simplify subsequent chromosomal integration of 23BD pathway genes using the thermostable Serine-recombinase assisted genome engineering (tSAGE) system (Ashok et al., 2024). Strains AG9463, AG9466, AG9468, and AG12932 were constructed using tSAGE (Ashok et al., 2024) by integrating plasmids pNA81, pNA85, pNA86, and pNA363, respectively, alongside the helper plasmid pNA42, which expresses the Y412MC61 serine recombinase.

#### Strain construction

*C. thermocellum* strains were constructed using tSAGE as described by (Ashok et al., 2024). Briefly, competent cells of *C. thermocellum* were co-transformed with a non-replicating integrating plasmid (pNA81, pNA83, pNA84, pNA85 and pNA86) and a non-replicating helper plasmid (pNA42). The integrating plasmids contained the *Geobacillus Y412MC61 attP* site, the genetic cargo to be delivered, and a *cat* gene that confers resistance to thiamphenicol (Table 1). The helper plasmid transiently expresses the *Geobacillus Y412MC61* serine recombinase from the Clo1313_2638 promoter. Following transformation via electroporation, the cells were recovered for approximately 18 hours in CTFUD at 50°C. The cells were then plated on CTFUD-Tm15 agar plates and incubated at 50°C. Colonies typically appeared after 2 to 3 days and were picked into CTFUD-Tm15 and incubated at 50°C. They were checked for plasmid integration by PCR and whole genome sequencing by Plasmidsaurus Inc. Complete genome sequences for each strain are included in Supporting Dataset D1.

**Table 1.** Plasmids used in this study.

| Name | Accession | Description |
| --- | --- | --- |
| pNA24 D | PX209995 | Plasmid replaces the *clo1313_2366* locus with a poly-*attB* cassette containing Bxb1, □BT1, □K38, □C1, RV, TG1, R4, |
| | | BL3, A118, MR11, SPβ, □370 and *Geobacillus* Y412MC61 *attB* sites through homologous recombination |
| pNA42 | PX209996 | tSAGE helper plasmid expressing the *Geobacillus* Y412MC61 serine recombinase |
| pNA81 | PX209997 | Plasmid carries *alsD* (*Bli03847*) and *gdh* (*Bli00828*) genes from *B. licheniformis* DSM 13 under the control of a strong *clo1313_1194* promoter. A 30 bp sequence, which includes the RBS from the *clo1313_1194* promoter, separates the genes. In the presence of the helper plasmid, the Y412MC61 *attP* site on the plasmid recombines with the corresponding *attB* site on the chromosome leading to the stable, single-copy insertion of the plasmid. |
| pNA83 | PX209998 | Plasmid carries *alsD* (*Bli03847*), *budC* (*Bli02066*) and *gdh* (*Bli00828*) genes from *B. licheniformis* DSM 13 under the control of a strong *clo1313_1194* promoter. A 30 bp sequence, which includes the RBS from the *clo1313_1194* promoter, separates the genes. In the presence of the helper plasmid, the Y412MC61 *attP* site on the plasmid recombines with the corresponding *attB* site on the chromosome leading to the stable, single-copy insertion of the plasmid. |
| pNA84 | PX209999 | Plasmid carries *alsD* (*Bli03847*), *budC* (*Bli02066*) and *gdh* (*Bli00828*) genes from *B. licheniformis* DSM 13 under the control of a strong *clo1313_1194* promoter. A 15 bp sequence that includes the canonical RBS separates the genes. In the presence of the helper plasmid, the Y412MC61 *attP* site on the plasmid recombines with the corresponding *attB* site on the chromosome leading to the stable, single-copy insertion of the plasmid. |
| pNA85 | PX210000 | Plasmid carries *alsD* (*Bli03847*) and *budC* (*Bli02066*) genes from *B. licheniformis* DSM 13 under the control of a strong *clo1313_1194* promoter. A 15 bp sequence that includes the canonical RBS separates the genes. In the presence of the |
|  |  | helper plasmid, the Y412MC61 *attP* site on the plasmid recombines with the corresponding *attB* site on the chromosome leading to the stable, single-copy insertion of the plasmid. |
| pNA86 | PX210001 | Plasmid carries *alsD* (*Bli03847*) and *budC* (*Bli02066*) genes from *B. licheniformis* DSM 13 under the control of a strong *clo1313_1194* promoter. A 30 bp sequence, which includes the RBS from the *clo1313_1194* promoter, separates the genes. In the presence of the helper plasmid, the Y412MC61 *attP* site on the plasmid recombines with the corresponding *attB* site on the chromosome leading to the stable, single-copy insertion of the plasmid. |

Note: Plasmids pNA85 and pNA86 have the same genetic cargo but have different ribosome binding sites between the two genes.

#### Plasmids

Plasmids were designed in-house and were built by Genscript Inc. Plasmids were maintained in and extracted from an *E. coli* strain lacking Dcm methylation (Guss et al., 2012) *E. coli* Top10 Δdcm (AG583) (Riley et al., 2019) grown in LB (Miller) medium, supplemented with 25 µg/mL chloramphenicol or 25 µg/mL of neomycin, with shaking at 37°C. The plasmids and strains used in this study are described in Table 2, and their sequences can be found in Supplemental File 1.

**Table 2.**
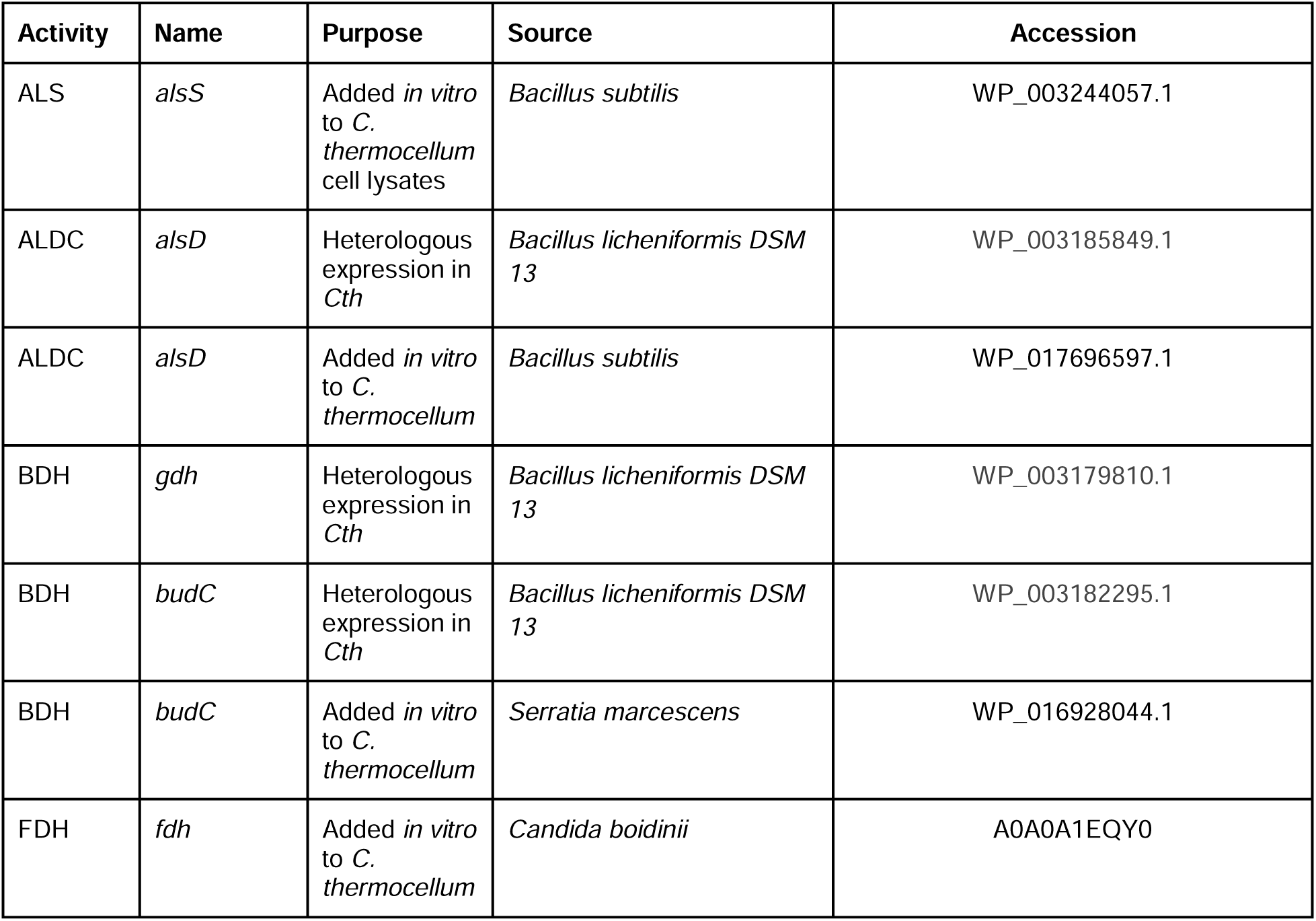
Genes used in this study.

#### Preparation of *E. coli* electrocompetent cells

An overnight culture of *E. coli* AG583 (Riley et al., 2019) culture was used to inoculate 50 mL of LB in a 250 mL flask, aiming for an optical density at 600 nm (OD_600_) of around 0.05. The culture was then incubated with shaking at 37 °C for approximately 2 to 3 hours until the OD600 reached between 0.5 and 0.7. Cells were pelleted by centrifugation at 7000 x g and 4°C, washed three times with ice-cold 10% glycerol and resuspended in 500 µL of the same buffer. Aliquots of 50 µL from the AG583 electrocompetent cells were either stored at -80 °C for later use or used immediately for plasmid transformation. Plasmids were electroporated into *E. coli* AG583 using a Bio-Rad Gene Pulser Xcell configured for exponential decay, with parameters set at 25 µF, 200 ohms, and 1800 volts, employing a 1 mm cuvette. After the electrical pulse, the cells in the cuvette were resuspended in 950 µL of SOC medium, followed by incubation at 30 °C for 3-4 hours. Subsequently, the cells were plated on LB media with appropriate antibiotics (25 µg/ml of chloramphenicol or 25 µg/ml of neomycin) and incubated overnight at 30 °C. Successful plasmid transformation was verified through PCR analysis of the colonies. Finally, plasmids were extracted from 50 mL liquid cultures (LB with suitable antibiotics) using a Zymo midi prep kit, following the low copy number plasmid extraction protocols.

#### Preparation of *C. thermocellum* electrocompetent cells

A -80 °C frozen stock of *C. thermocellum* was used to inoculate 5 mL of CTFUD rich medium and was grown at 50 °C inside the Coy anaerobic chamber. This culture was then used to inoculate 500 mL of CTFUD, which was grown overnight at 50 °C anaerobically to an OD_600_ between 0.5 and 0.7. The culture was chilled on ice, pelleted by centrifugation at 7000 X g for 10 minutes at 19°C, and washed with ice-cold electroporation buffer (250 mM sucrose and 10% glycerol) three times. After each centrifugation, the supernatant was carefully poured off, and fresh electroporation buffer was added without disturbing the pellet. No resuspension of the pellet occurred during the washes. After the third wash, 100 µL of fresh electroporation buffer was added, and the cells were resuspended by pipetting up and down, resulting in a total volume of approximately 500 µL of competent cells. The cells were then immediately used for plasmid transformation or frozen at - 80 °C for later use. 50 µL of competent cells and one µg of each plasmid (extracted from AG583) were combined in a 1 mm electroporation cuvette. The Bio-Rad Gene Pulser was set to a square wave pulse of 1000V for 1.5 msec. Following electroporation, the cells were resuspended in 5 mL of CTFUD and recovered overnight at 50 °C inside the anaerobic chamber. The recovered cells were then plated on CTFUD-Tm15 agar plates.

#### Cell pellet preparation for enzyme assays

For measurement of enzyme activity, cell pellets were prepared by growing strains in MTC-5 chemically defined medium with 5 g/L cellobiose in 15 mL tubes at 55 °C under anaerobic conditions. At OD_600_ ∼ 0.4, cells were harvested by centrifugation. The resulting cell pellets were washed with 50 mM HEPES buffer and suspended in 250 μL final volume. Cells were lysed by the addition of 1 μL of Ready-Lyse lysozyme solution (Epicenter, WI, USA) and incubated at room temperature for 20 min. Then, 1 μL of DNase I solution (Thermo Scientific, MA, USA) was added, and lysate was incubated for an additional 20 min at room temperature. The lysate was centrifuged under anaerobic conditions at 12,100 RCF for 5 min. The supernatant was harvested, and its protein concentration was determined using the BCA assay (G-Biosciences, Solon, OH). This resulting supernatant is the cell lysate that was used in subsequent enzyme assays.

#### Enzyme preparations

Als, Aldh, Fdh, and Bdh enzymes for *in vitro* assays with lysates were produced following previously described protocols (Jilani et al., 2024).

#### Enzyme activity assays

All enzyme activity assays were performed in HP-Agilent (Model 8453) UV-Vis diode array spectrophotometer at 25 °C under following enzyme specific conditions:

#### Acetolactate synthase (ALS) enzyme

The ALS assay was performed in 1 mL reaction mix consisting of 50 mM HEPES and 66 mM sodium pyruvate. The reaction was started by addition of cell lysate. And reaction progress was observed by monitoring the change in absorbance at 320 nm corresponding to the absorbance by sodium pyruvate.

#### Acetolactate decarboxylase (ALDC) enzyme

ALDC specific activity was determined via coupled assay as described before (Jilani et al., 2024). Briefly, the assay was performed in 1 mL reaction mix consisting of 50 mM HEPES, 66 mM sodium-pyruvate, 0.02 mg Als enzyme, 5 mM NADH, 0.19 mg Bdh enzyme. The reaction was started by addition of cell lysate and Aldc specific activity was determined by calculating the slope of the decrease in 390 nm signal corresponding to the reduction of NADH in the reaction mix. A wavelength of 390 nm was used instead of the more commonly-used 340 nm to reduce interference from pyruvate (Jilani et al., 2024).

#### 23BD dehydrogenase (BDH) enzyme

The BDH assay was conducted in a 1 mL reaction mix consisting of 50 mM HEPES, 15 mM acetoin and 0.2 mM NADH. The reaction was initiated by the addition of cell lysate to the reaction mix. The Bdh specific activity was determined by calculating the slope of the decrease in 340 nm signal corresponding to a reduction in NADH in the reaction mix.

#### Cell lysate experiments

For cell lysate experiments, cells were prepared as described for other enzyme assays, but washed with a “cytoplasmic buffer” consisting of 50 mM HEPES, 5 mM MgCl_2_, 60 mM KCl, 50 mM NaHCO_3_, 5 mM DTT, and 5 mM NH_4_Cl (Jilani et al., 2024) and resuspended in 250 μL final volume. Cell lysis was also performed as described above. The concentration of protein in the cell extract was 3 mg/mL. The initial NAD^+^ concentration was 1 mM. The initial formate concentration was 100 mM. The initial pyruvate concentration was 60 mM. In cases where purified enzymes were added to cell lysate, the following concentrations were used (mg/mL): acetolactate synthase (Als), 0.02; acetolactate decarboxylase (Aldc), 0.15; 2,3-butanediol dehydrogenase (Bdh), 0.19; and formate dehydrogenase (Fdh), 0.8. The experiment was conducted in a 50 µL reaction mix in a 200 µL reaction tube with reduced volume cap at 37 °C under anaerobic conditions, similar to our previous work with this system (Jilani et al., 2024). The metabolites were quantified by HPLC after 24 h incubation.

#### Quantification of fermentation products

Screening of all generated mutant strains was performed in 15 mL tubes containing 5 mL of MTC-5 medium and 5 g/L cellobiose. Fermentation cultures were incubated at 55 °C for 24 hours. Metabolites were quantified by high-pressure liquid chromatography (HPLC). Samples were treated with 2.5 % trichloroacetic acid (TCA) to precipitate proteins. 0.5% sulfuric acid was added to the samples before analysis. Metabolites were measured using a Shimadzu-HPLC (Model 2030 C-3D) equipped with an Aminex HPX 87H (300 × 7.8mm) column equipped with RI and PDA detectors. Column and detector temperature was maintained at 60 °C and detector at 40 °C, respectively. 2.5 mM sulfuric acid was used as the mobile phase with a flow rate of 0.6 mL/min. We observed that the SS and RR isoforms of 23BD coelute, which makes it difficult to differentiate between the two isoforms. However, literature suggests (Ge et al., 2016) that the 23BD gene from *Bacillus licheniformis* catalyzes formation of RR isoform of 23BD. Standards of each analyte were freshly prepared before analysis. Limits of detection for acetoin, m23BD and SS/RR23BD were 0.062, 0.062 and 0.25 mM, respectively.

## Results and Discussion

### Identification of candidate genes and gene expression elements

To establish a functional 23BD pathway in *C. thermocellum*, we leveraged previous work on *Bacillus licheniformis* DSM 13. This strain is a natural 23BD producer that thrives between 37 °C and 55 °C (Dong et al., 2018) deletion of the *gdh* and *budC* genes in *B. licheniformis* causes an accumulation of acetoin, a key pathway intermediate (Lü et al., 2020).Using this information, we designed gene constructs to express the *B. licheniformis* genes *alsD*, *gdh*, and *budC* in *C. thermocellum* (Figure 2). Based on findings from Olson et al in 2015 (Olson et al., 2015) that identified the P_Clo1313_1194_ promoter is highly expressed in *C. thermocellum* (Olson et al., 2015) and was recently found to be the strongest promoter tested when expressed from the *C. thermocellum* chromosome (Ashok et al., 2024). Therefore, we created three artificial operons: one with P_Clo1313_1194_ driving *alsD* and *gdh*, another with *alsD* and *budC*, and a third with all three genes. These constructs were then integrated into the chromosome of *C. thermocellum* strain AG8235 using tSAGE and verified with whole-genome sequencing (Supporting Dataset D1)(Table 3).

**Figure 2.**
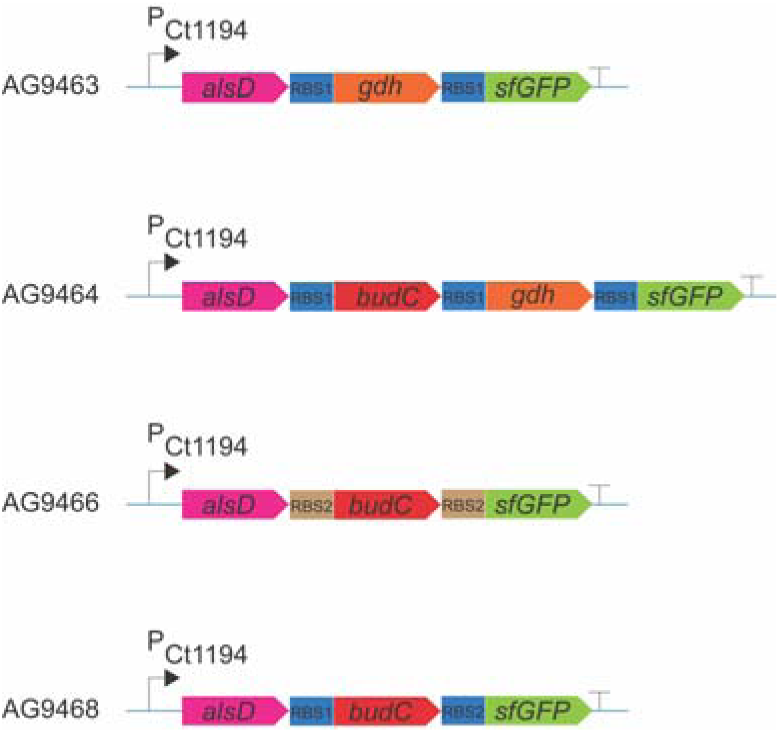
Cassette design for 23BD production pathways. P_Ct1194_, *C. thermocellum Clo1313_1194* promoter; *alsD*, *B. licheniformis* acetolactate decarboxylase; *gdh*, *B. licheniformis* RR-23BD dehydrogenase gene; *budC*, *B. licheniformis* meso-23BD dehydrogenase gene; *sfGFP*, superfolder green fluorescence protein gene; RBS1, ribosome binding site and spacer from the Clo1313_1194 gene (cttaattatcattaaagggggaaaaaaact); RBS2, synthetic ribosome binding site and spacer (aggaggaatagtctc); T, terminators L3S1P13 (Chen et al., 2013) and ECK120010850 (Vasudevan et al., 2019).

**Table 3.** Strains used in this study, the plasmids that facilitated their creation, the methods employed in the construction process, and the genetic material carried by each integration plasmid.

| Organism | Strain ID | Parent ID | ALDC | BDH | Cassette | Construction notes |
| --- | --- | --- | --- | --- | --- | --- |
| *E. coli* | AG9163 |  |  |  |  | *E. coli* TOP10 derived strain with a di-methylation system, for improved transformation of *thermocellum* |
| *C. thermocellum* | LL1299 |  |  |  |  | (Tian et al., 2019) |
|  | AG8235 | LL1299 |  |  | None | (Ashok et al., 2024) |
| | AG9463 | AG8235 | *alsD* | *gdh* | $P_{1194}\text{-alsD-gdh}$ | Plasmid pNA81 integrated via tSAGE |
| | AG9464 | AG8235 | *alsD* | *gdh, budC* | $P_{1194}\text{-alsD-budC-gdh}$ | Plasmid pNA83 integrated via tSAGE |
| | AG9465 | AG8235 | *alsD* | *gdh, budC* | $P_{1194}\text{-alsD-budC-gdh}$ | Plasmid pNA84 integrated via tSAGE |
| | AG9466 | AG8235 | *alsD* | *budC* | $P_{1194}\text{-alsD-budC}$ | Plasmid pNA85 integrated via tSAGE |
| | AG9468 | AG8235 | *alsD* | *budC* | $P_{1194}\text{-alsD-budC}$ | Plasmid pNA86 integrated via tSAGE |

### Initial Characterization of Engineered Strains

Initially, strains were screened for production of 23BD in MTC-5 chemically defined medium with 5 g/L cellobiose as carbon source (Figure 3A). Strain AG9463 produced 4.1 mM 23BD, while 52 and 60-fold lower levels were observed in strains AG9466 and AG9468, respectively. The theoretical maximum yield from 5 g/L cellobiose (14.7 mM) is 29.4 mM 23BD, so 4.1 mM 23BD represents 14% yield. No 23BD was detected in the fermentation broth of strain AG9464.

**Figure 3.**
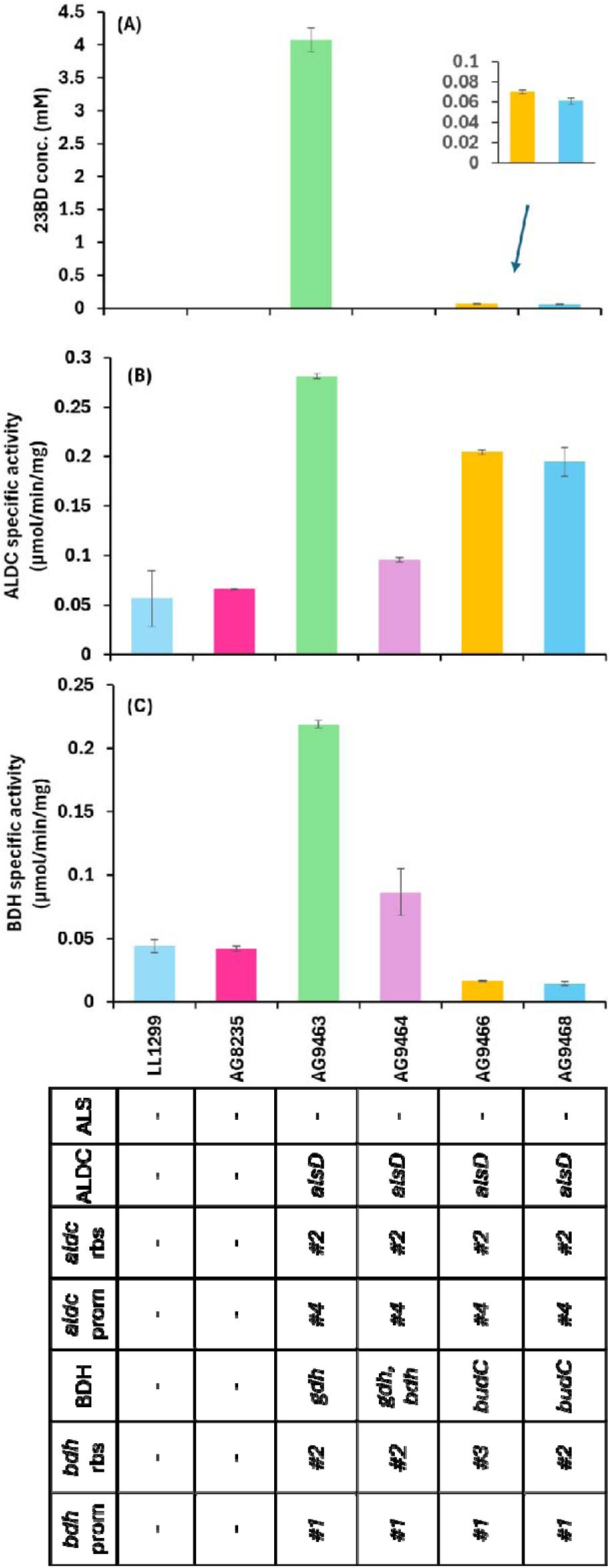
Analysis of 23BD production in engineered strains of *C. thermocellum*. All strains were cultured in MTC-5 medium containing 5 g/L cellobiose as carbon source. (A) 23BD titers detected in the fermentation broth are the sum of both SS/RR 23BD and meso-23BD. (B) ALDC enzyme activity. (C) BDH enzyme activity. For both ALDC and BDH, assay conditions are described in the materials and methods section. Error bars represent the range of data, n=2 biological replicates. Individual replicate data, and separate quantification of 23BD isomers, is available in Supporting Dataset D2. *AG8235 is the parent strain used to construct all the other strains listed in this table Δhpt Δ0478 Δ2366::polyattB. #1 – p1194, promoter from the Clo1313_1194 gene #2 – aggggga ribosome binding site sequence #3 – aggagga ribosome binding site sequence #4 – P_Clo1313_1194 (1^st^ gene in the operon)

To further investigate whether the cloned genes are being translated into functional enzymes, we measured the enzyme activity of respective cloned gene(s) in each strain in crude cell lysates. Although low levels of ALDC activity are observed in the parent strains, expression of the *alsD* gene increased ALDC activity 4 to 6-fold (Figure 3B). The one exception is strain AG9464. However, the lower ALDC activity in this strain may be due to a spontaneous mutation that we later found in the P_Ct1194_ promoter region consisting of a duplication of the 21 bp sequence 5’ - TAATTTATAATATAATTTATT - 3’ 64 bp upstream of the *alsD* start codon (Supporting Text T1).

BDH enzyme activity was also measured for the two putative *bdh* genes: *budC* and *gdh* (Figure 3C). The maximum values of BDH activity were observed in strain AG9463, followed by strain AG9464. Between the two, *gdh* appears to confer higher activity, and this was associated with increased 23BD production. The *budC* genes were tested with two different ribosome binding site (RBS) sequences, but there was no difference between these in terms of either 23BD production or BDH activity. Increased BDH activity was a stronger predictor of final 23BD titer, compared to ALDC activity, suggesting that BDH has more control of pathway flux. All of the observed BDH activity was NADH-linked. No activity was observed with NADPH as a cofactor (data not shown).

No activity of the endogenous Als enzyme was detected in our enzyme assay; however, this activity must be present in vivo due to the observed production of 23BD and the native ability to synthesize valine.

### Cell-free analysis of pathway function

The presence of 23BD production in strain AG9463 demonstrated that a functional pathway was present, however the product yield was relatively low. We hypothesized that the yield could be limited by a lack of a particular pathway enzyme or problems with redox balance. We therefore used a cell-free systems biology approach (Jilani et al., 2024) to diagnose this problem using cell lysates.

We previously developed a four-enzyme system for in vitro conversion of pyruvate and formate to 23BD. In this system, the combination of formate and the formate dehydrogenase (FDH) enzyme serves as a cofactor recycling system to regenerate the NADH used by the BDH reaction (Jilani et al., 2024). We focused our characterization on strain AG9463 due to it having the highest levels of BDH and ALDH activities and demonstrated 23BD production.

To understand the pyruvate to 23BD pathway in the parent strain and in strain AG9463, we added purified enzymes in various combinations to the cell lysate to identify factors limiting 23BD production (Figure 4). Cell extract from the parent strain (LL1299), when supplemented with Fdh to generate NADH and all three pathway enzymes (Txt1), converts pyruvate stoichiometrically to 23BD. When BDH is left out of the reaction, acetoin accumulates instead of 23BD, indicating that BDH activity is lacking in the parent strain (Txt2). When all three pathway enzymes are left out (Txt3), a small amount of acetoin is made, as is some ethanol, indicating that functional ALS and ALDC activities exist in the parent, but that they do not support high flux. (Note: although acetoin can be produced non-enzymatically from diacetyl, this reaction is only thought to proceed in the presence of oxygen, and our cell-free reactions were performed under anaerobic conditions). This is further supported by the reaction in which BDH is the only pathway enzyme added (Txt4), where some of the low level acetoin is instead converted to 23BD.

**Figure 4.**
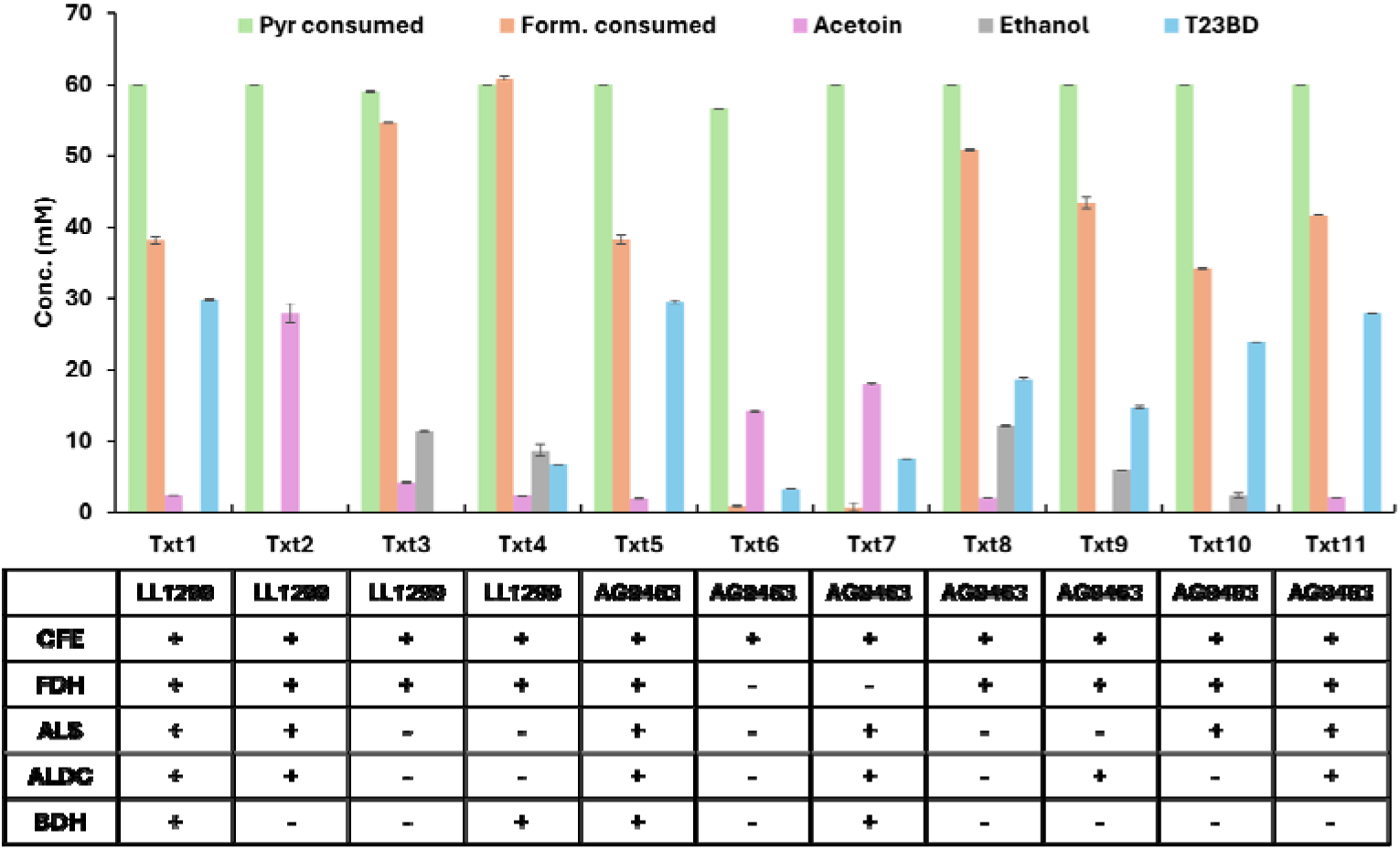
Analysis of mutant strains via cell free lysate reactions. Lysates of strains LL1299 and AG9463 were analyzed by multiple permutations and combinations of the heterologous purified enzymes involved in conversion of pyruvate to 23BD. Each treatment is labeled with a unique ID (Txt1, Txt2, etc.). The composition of each lysate is described below the corresponding treatment. The initial concentration of formate was 100 mM. The initial concentration of pyruvate was 60 mM. Cell-free reactions were run for 24 h and quantified by HPLC. Additional details are provided in the materials and methods. Error bars represent the range of the data, n=2. Individual replicate data, and separate quantification of 23BD isomers, are provided in Supporting Dataset D2.

We next used the cell-free system to characterize strain AG9463. Cell extract from this strain, in the presence of FDH as a source of NADH and with all pathway proteins added (Txt5), was able to catalyze the conversion of pyruvate to 23BD, confirming that the cell lysate does not inhibit activity of the exogenous added enzymes. When FDH is left out, either with or without added pathway enzymes (Txt6 and Txt7), the extract primarily produces acetoin, consistent with the fact that acetoin reduction to 23BD requires NADH.

With the addition of FDH to provide NADH but no exogenous enzymes (Txt8), substantial flux to 23BD is observed, indicating that a functional pathway is present in the strain. However, the yield of 23BD was not stoichiometric, indicating that further pathway optimization is possible. The further addition of ALDC did not improve production (Txt9), but the addition of ALS did further improve 23BD production (Txt10). The addition of both ALS and ALDC increased 23BD production to near-stoichiometric conversion (Txt11), similar to the reaction with all components added (Txt5). Therefore, increasing ALS activity is a promising target for future metabolic engineering, and increasing ALDC activity may also be beneficial. We also confirmed that BDH expression is sufficient for high yield 23BD production, since addition of purified BDH enzyme to the AG9463 lysate (which expresses a heterologous Bdh enzyme) did not affect 23BD yield (compare Txt5 vs Txt11).

Further analysis of the data from the cell-free system allowed us to determine the relative importance of different interventions for increasing 23BD production (Figure 5). The effect of the NADH recycling system (consisting of formate plus the Fdh enzyme) was the most important intervention, indicating the importance of redox balance for driving flux through the 23BD pathway. The addition of the Als enzyme was the next most important intervention, followed by addition of Aldc. There was no benefit to addition of the Bdh enzyme (and in some cases, even a slight decrease in yield).

**Figure 5.**
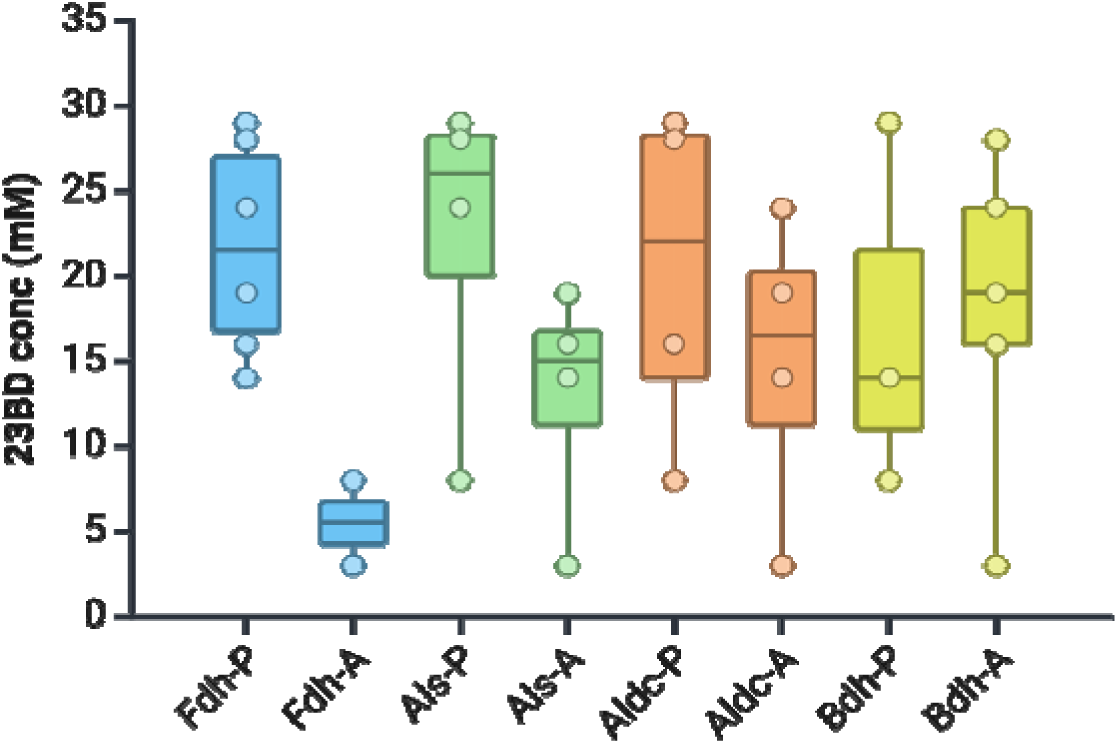
Relative contribution of each heterologous purified enzyme in determining observed 23BD titers in cell lysates of strain AG9463. The letters “P” and “A” in the axis labels represent either presence or absence, respectively, of each enzyme in the lysate treatment. Boxes are colored by enzyme and describe the 25th-75th percentile of the data. Dots represent each data point in respective treatment.

### Addition of acetate improves yield of 23BD

Since sugar fermentative metabolism is characterized by excess NADH which leads to cellular redox imbalance, we explored methods to reduce or minimize the imbalance and improve titers of 23BD. We first tested the addition of acetaldehyde, which can be reduced to ethanol by the alcohol dehydrogenase (ADH) reaction, thereby acting as an electron sink; however, we observed minimal effects on 23BD production and substantial growth inhibition (Supporting Figure S1). We suspect that aldehyde toxicity was greater than any potential redox effect offered by acetaldehyde to serve as a sink of excess electrons.

We also tested the addition of acetate to the fermentation (Figure 6). In *C. thermocellum*, acetate is normally a fermentation product, and similar to 23BD, acetate production results in the production of excess reducing equivalents that need to be oxidized, for instance by production of a more-reduced compound like formate or H_2_. We hypothesized that addition of acetate to the growth medium might decrease acetate biosynthesis, thereby decreasing the overall redox imbalance. We therefore evaluated the fermentation behavior of the 23BD-producing strain (AG9463) in presence of low (14.6 mM) and high (58.4 mM) cellobiose loadings in MTC-5 defined medium. The low cellobiose condition was designed to ensure complete cellobiose consumption. This high cellobiose condition was designed to ensure that some cellobiose remained (i.e non-limiting concentration) at the end of the fermentation.

**Figure 6.**
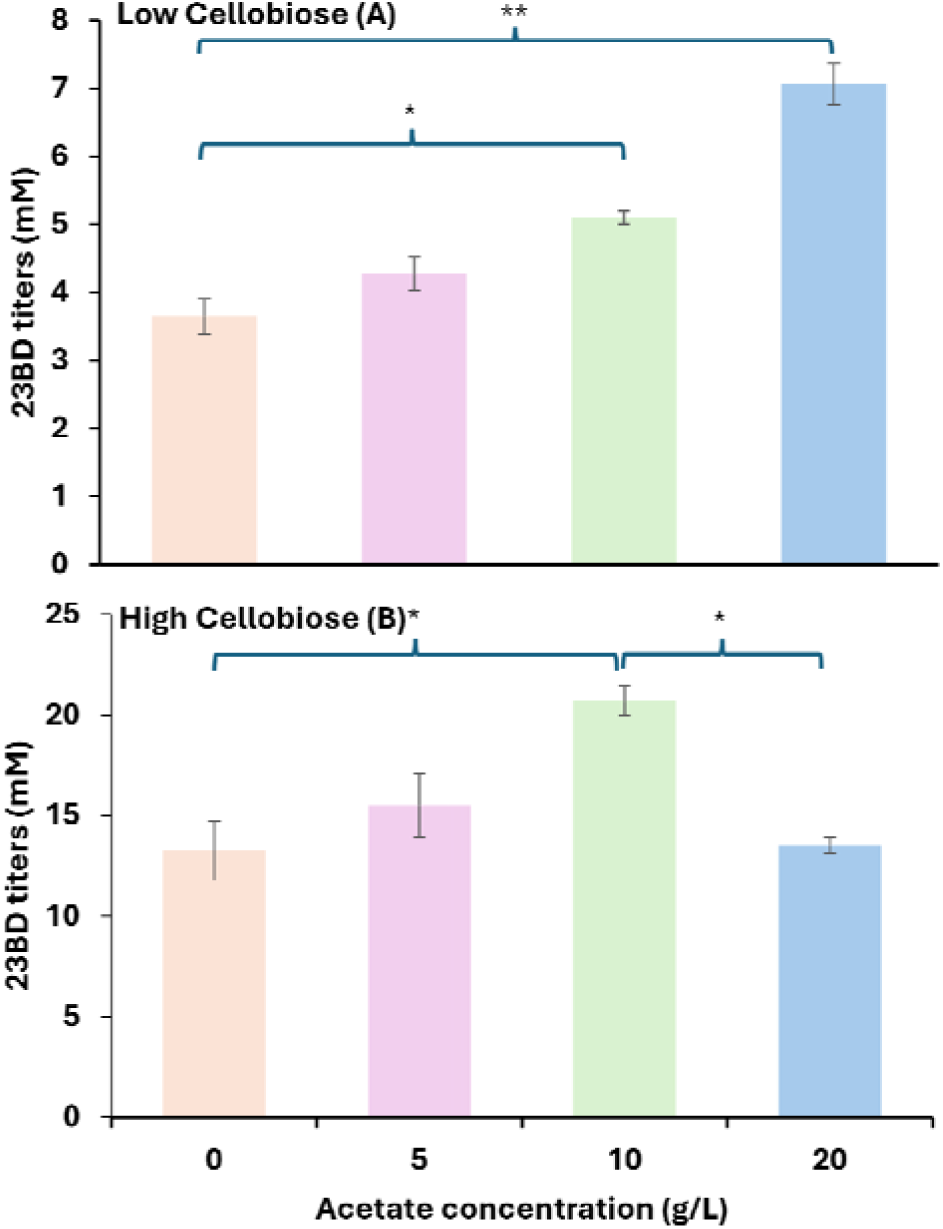
Influence of acetate supplementation on titers of 2,3-BD in *C. thermocellum* AG9463. Cultures were cultivated in 50 mL MTC-5 media with either 5 (Low) or 20 g/L (High) cellobiose concentration, respectively. Acetate concentrations of 5, 10 and 20 g/L correspond to 61, 122 and 244 mM, respectively. Error bars represent the range of the data, n=2 biological replicates. Tests were performed using R 4.2.2 and One-way ANOVA with Tukey-multiple comparison test. * represents p-value=0.03; ** represents p-value=0.001. Complete fermentation data is available in Supporting Dataset D2.

Increasing the substrate concentration increased 23BD titers from 4 mM to 13 mM, with the yield decreasing slightly at increased substrate loading. For both cellobiose concentrations, we tested various acetate concentrations (5, 10, and 20 g/L). The addition of acetate increased 23BD yield, with the largest yield increases being observed at 10-20 g/L (depending on the initial substrate concentration) (Figure 6). We observed a negative interaction between the 20 g/L acetate and high (20 g/L) cellobiose loadings for 23BD production.

We also tested the influence of variation in pH, in the presence of acetate, on metabolite distribution (Supplementary Figure S3). At pH 6 no growth was observed in either strain LL1299 or strain AG9463. As expected, no 23BD was detected from cultures of strain LL1299.

Interestingly, we observed an increase in 23BD titers with an increase in pH of the medium for strain AG9463. We predict that with an increase in pH of the medium, the dissociated acetate anion is able to modify cellular redox homeostasis and redistribute carbon flow away from ethanol towards production of 23BD. We do not know if acetate inhibits Pfor or AdhE enzymes involved in conversion of pyruvate to ethanol. More investigation is necessary to arrive at a definitive explanation behind our observation.

One of the initial goals of this project was to demonstrate 23BD production from cellulose, and this is why we chose *C. thermocellum* as a host organism. *C. thermocellum* has the remarkable ability to ferment crystalline cellulose as well as (or in some cases even better than) soluble sugars such as cellobiose (Holwerda et al., 2013). Therefore, we wanted to make sure that ability had not been lost during our work to engineer the organism to produce 23BD. Having tested the ability of *C. thermocellum* AG9463 to utilize cellobiose, we tested its ability to convert crystalline cellulosic carbon to 23BD (Figure 7). Prior work has shown that small amounts of 23BD (0.42 g/L or 4.66 mM) are produced when *C. thermocellum* is grown on very high (>100 g/L) substrate concentrations (Holwerda et al., 2014). Therefore, we grew the strain on lower substrate concentrations (20 g/L, equivalent to 123 mM glucose) to avoid the confounding effects of overflow metabolism.

**Figure 7:**
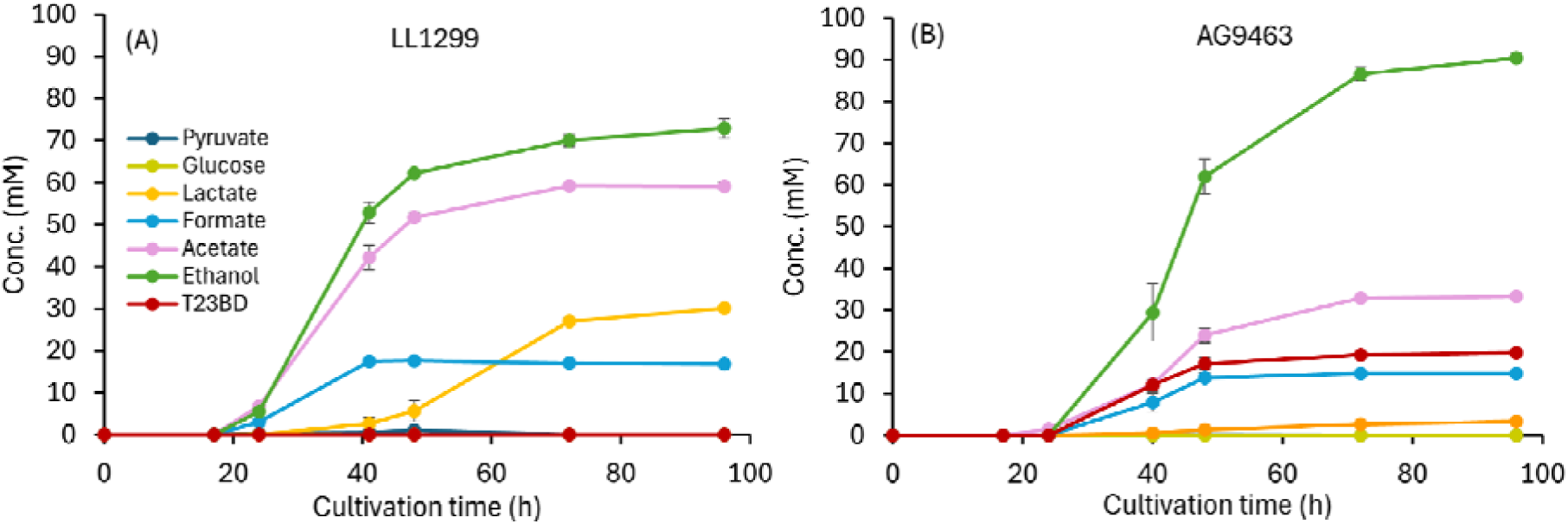
Utilization of Avicel microcrystalline cellulose as a carbon source. Strains LL1299 (A) and AG9463 (B) were cultivated in presence of 20 g/L Avicel in 50 mL MTC-5 medium (approx. 123 mM glucan equivalents). Cultivation was performed in serum bottles at 55 □ with shaking at 180 RPM. Samples were extracted at indicated time points in an anaerobic chamber. Error bars represent the range of the data, n=2 biological replicates.

In the absence of heterologous expression of 23BD pathway genes (strain LL1299), 23BD production was not observed; however, the strain expressing *gdh* and *alsD* (strain AG9463) showed high levels of 23BD production (19.7 mM, Figure 7). Interestingly, we observed a remarkable redistribution of carbon metabolism in AG9463 as compared to the parent LL1299. At 96 h, strain AG9463 produced almost 10-fold less lactate than the parent strain LL1299 (30.1 mM vs 3.3 mM). Lactate production is redox-neutral. However, its production is generally thought to indicate an accumulation of glycolytic intermediates such as fructose-1,6-bisphoshpate (FBP) (Holwerda et al., 2020; Rydzak et al., 2015). Thus, the decreased lactate production in strain AG9463 suggests that 23BD production may serve as a drain for pyruvate in central metabolism that prevents accumulation of FBP.

A reduction in the concentration of acetate was also observed, decreasing from 59.2 mM in the control strain to 33.2 mM in AG9463. A corresponding 24% increase in the titer of ethanol was observed in strain AG9463 to 90.4 mM from 72.9 mM observed for the parent LL1299 strain.Both strains produced approximately equal amounts of total products (∼165 mM on a C3 basis, or 83 mM on a glucan (i.e. C6) basis). Of the 123 mM glucan equivalents present in the substrate, about two thirds (∼83 mM) was converted to detectable fermentation products, a typical value for this organism (Ellis et al., 2012; Holwerda et al., 2020, 2012). Thus the 19.7 mM final 23BD titer represents a metabolic yield of 24% (23BD per total products). On an available electron basis (Villadsen et al., 2011), the total fermentation products from strain LL1299 resulted in 1726 mM available electrons vs. 1839 mM from strain AG9463, a 6.6% increase. Since the carbon balances are approximately equal, but there is an increased amount of available electrons in strain AG9463, increased 23BD production cannot be completely explained by the decreased acetate and increased ethanol production. The most likely source of these additional electrons is from reduced hydrogen production, although this was not measured.

Since conversion of glucans to 23BD is not redox balanced, the maximum theoretical yield of 23BD in *C. thermocellum* depends on assumptions about how the strain eliminates excess reducing equivalents generated in glycolysis. One way to do this is via a combination of ethanol and hydrogen production. If the cell uses a bifurcating hydrogenase (Buckel and Thauer, 2013) (Fd_red_^2-^ + NADH + 3H^+^ ⇔ Fd_ox_ + NAD^+^ + 2H_2_) to consume electrons from Fd during ethanol production, then the net reaction during ethanol production is ½ glucan + 2 NADH = ethanol + 2 H_2_ + 2 NAD^+^. Thus, ¼ glucan is required to eliminate each excess reducing equivalent (e.g., NADH) that is generated during 23BD production, resulting in a maximum theoretical yield of 23BD from glucose of (1/1.25 = 0.8). Thus our 24% metabolic yield represents about 30% of the theoretical maximum yield for *C. thermocellum* when redox balance constraints are taken into account.

## Conclusions

We demonstrated a proof of concept functional expression of a thermophilic 23BD production pathway in *C. thermocellum*, resulting in a yield up to 24% of the theoretical maximum. We further demonstrated the utility of cell-free systems for troubleshooting metabolic pathway engineering in non-model organisms, such as *C. thermocellum*.

Cell-free supplementation demonstrated that added BDH activity was required for 23BD production in the parent strain of *C. thermocellum*, demonstrating that *C. thermocellum* contains native genes that allow ALS and ALDC activity (Figure 4). Two promising candidate genes that could catalyze this ALS activity (Clo1313_0099 and Clo1313_0305), which are predicted to catalyze the same reaction for branched chain amino acid biosynthesis. For ALDC activity, on the the hand, a BLAST search of the *C. thermocellum* genome did not reveal any likely candidates for genes for this reaction, so the native route to acetoin remains unknown.

We evaluated the three enzymatic steps required to get from central energy metabolism to 23BD. ALS activity: converting two pyruvate into one acetolactate, as demonstrated by the high native flux to valine biosynthesis in *C. thermocellum* and by our cell-free assays. ALDC activity: converting acetolactate into acetoin, is also present natively in *C. thermocellum*, but it is limiting when BDH is abundant. Finally, BDH activity: converting acetoin into 23BD, is undetectable in *C. thermocellum* natively, as evidenced by the lack of 23BD production *in vivo* and *in vitro* without the addition of a BDH.

Further engineering is needed to increase flux to 23BD. Gene deletions may limit the production of other products, but they could also complicate challenges with redox balancing. Furthermore, an important area for future investigation is the effect of the cofactor specificity of the BDH reaction. In this work, we used an enzyme with NADH-linked activity. Although glycolysis typically results in generation of NADH, the atypical glycolysis of *C. thermocellum* allows generation of either NADH or NADPH, depending on the relative flux between the pyruvate phosphate dikinase reaction and the malate shunt (Olson et al., 2017; Zhou et al., 2013).

## Acknowledgements

None.

