## Supplementary data for "Engineering *Clostridium thermocellum* for production of 2,3-butanediol from cellulose"

### Supporting Information

**Supporting Dataset D1**. A text file with whole-genome resequencing data from strains LL1299, AG8235, AG9463, AG9464, AG9466, and AG9468 in Genbank format.

**Supporting Dataset D2.** Excel file containing individual replicate data used in the generation of figures in the manuscript. Each tab contains data for a separate figure.

**Supporting Text T1.** Mutation in the P_Ct1194_ promoter region in strain AMG9464. The repeated sequence is shown in bold. The first repeat is in black text, the second repeat is in red text. The start codon of the *alsD* gene is underlined.

5’ - TTAATATGCCGACCACGTTGCAATTCCCGTCAAATAATGCATTTTGCAGCCGACGAAACAGGCAAGATAACTGTATTGGCTATAAATGTTTCAGGCAGCGGTATATTTTGCCTCCCGGTAAAATTAATACAATAAGCTAAAAAACTGACGTAGGATAAGCAAAACGGCGCAATTTGAGTTGTAACGTAATATTTTCACTAAAAATAGTAATTATTTCATGTTGTTTTTTTTTAGAT**TAATTTATAATATAATTTATTTAATTTATAATATAATTTATT**GTATAAGCAATATCTTAATTATCATTAAAGGGGGAAAAAAACTATG - 3’

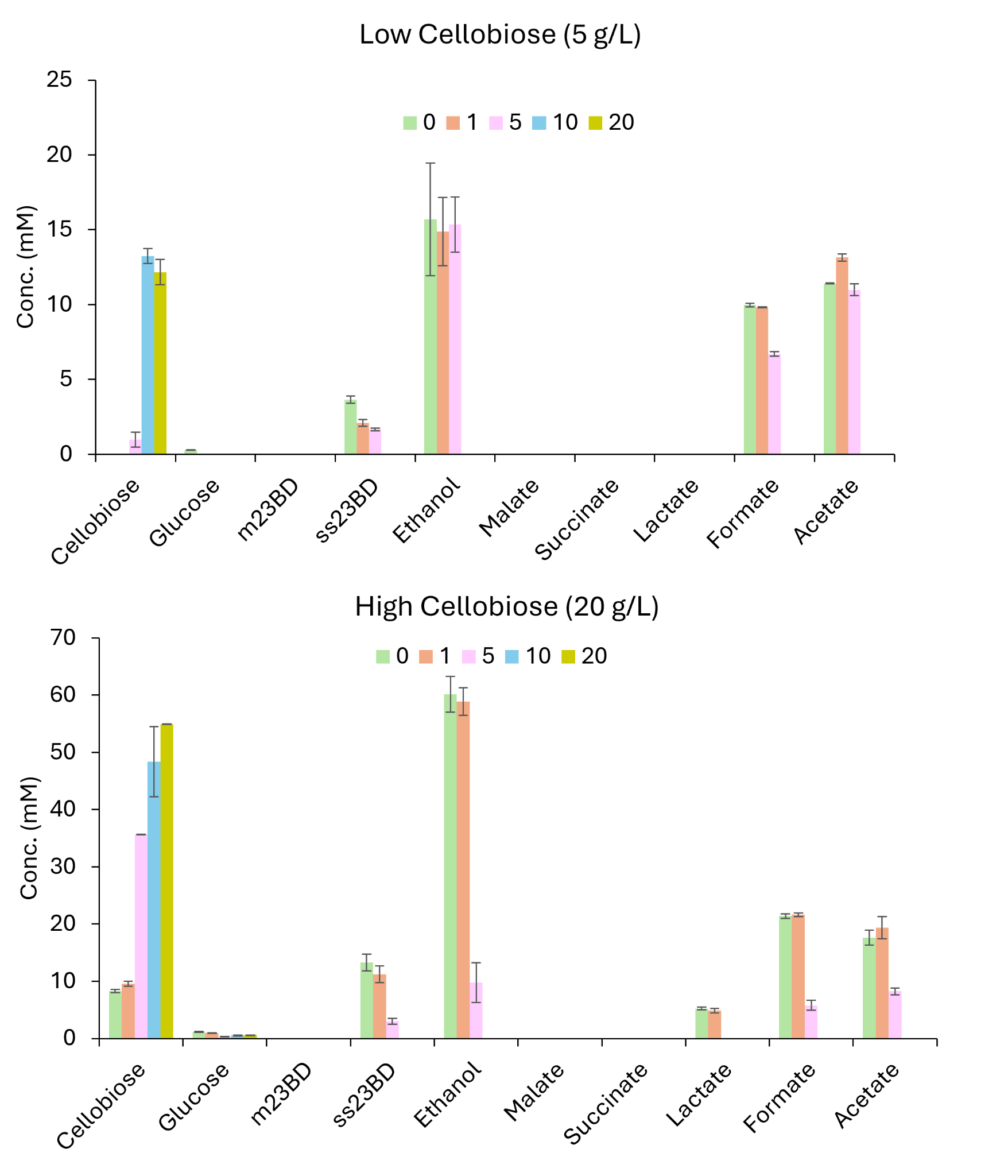

**Supporting Figure S1.** Influence of acetaldehyde (added as a possible external electron acceptor) on titers of 23BD in *C. thermocellum* strain AG9463. Cultures were cultivated in 50 mL MTC-5 media with either low (5 g/L) or high (20 g/L) initial cellobiose concentrations. Acetaldehyde concentrations of 1, 5, 10 and 20 g/L correspond to 23, 114, 227 and 454 mM, respectively. Error bars represent the range of the data, n=2 biological replicates.

**Supporting Figure S2**. Chromatogram showing HPLC traces for key metabolites (ss/rr23BD, rr23BD, meso23BD, acetoin, pyruvate, etc.

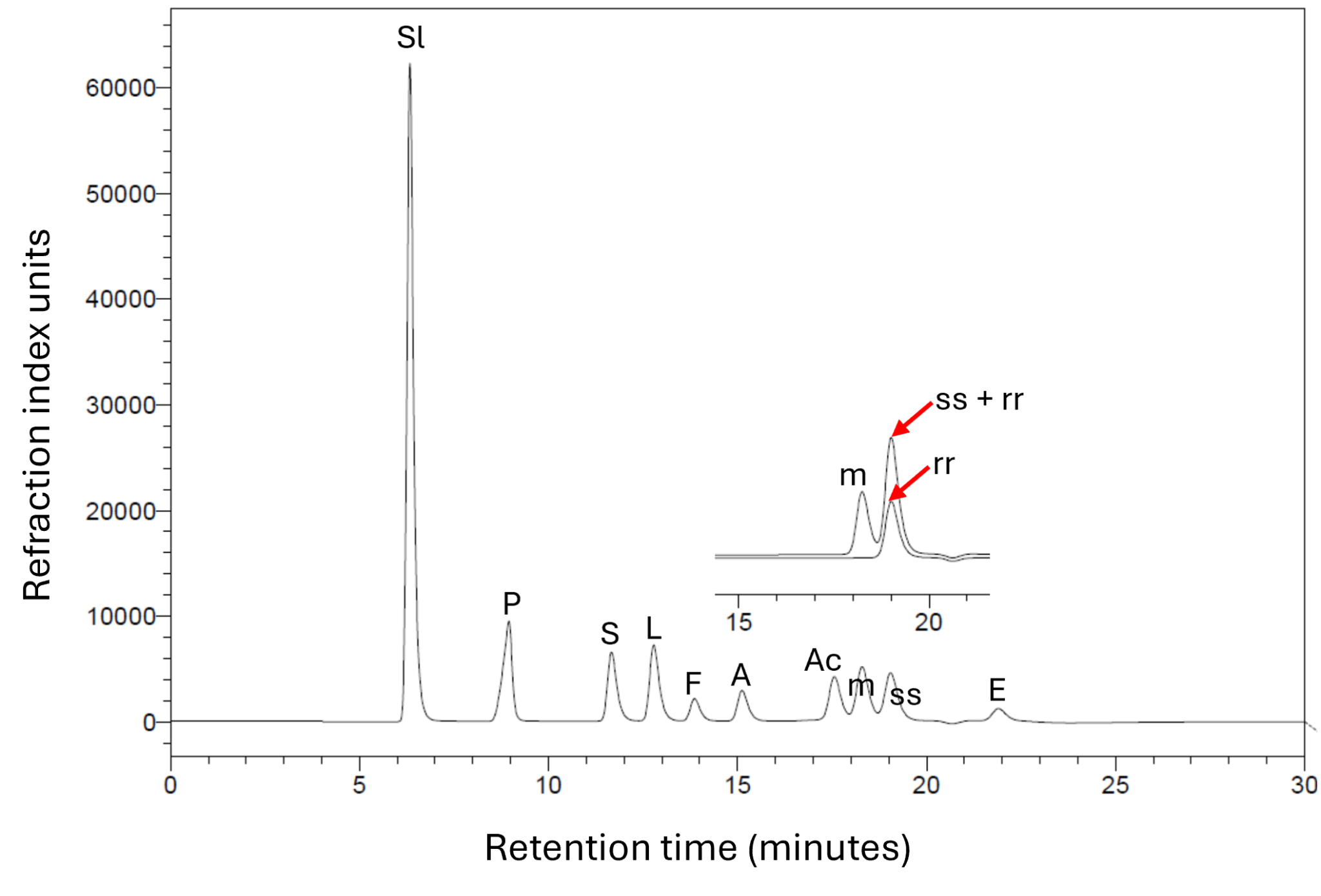

**Supporting Figure S2**: HPLC traces for key metabolites.both ss and rr isoforms of 23BD coelute at the same retention time which leads to an additive increase in the peak height. Sl - sulfuric acid; P - pyruvate; S - succinate; L - lactate; F - formate; A - acetate; Ac - acetoin; m - meso-2,3-butanediol; SS-2.3-butanediol; rr - RR-2,3-butanediol; E - ethanol.

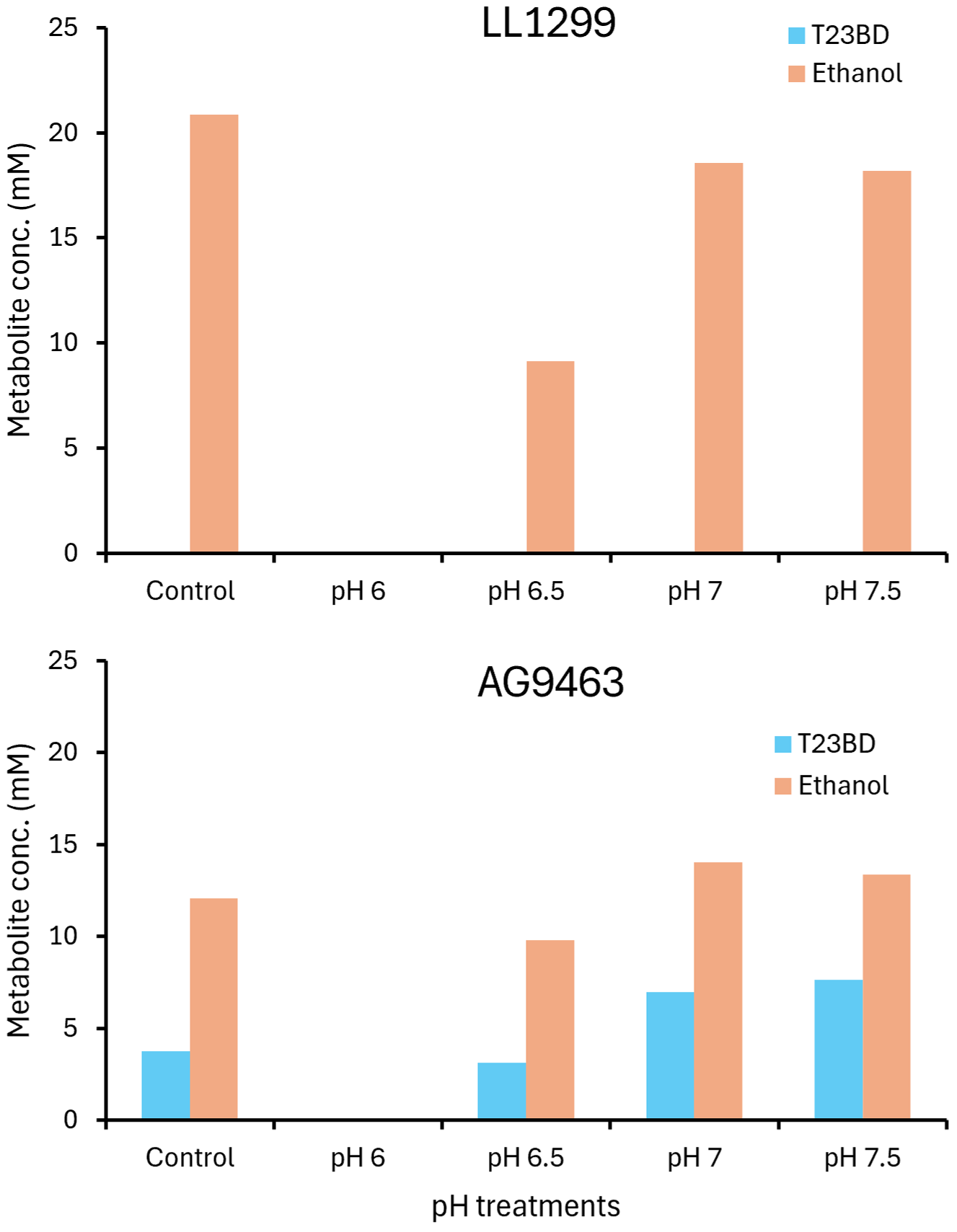

**Supporting Figure S3**: Influence of amending acetate (10 g/L) to culture medium on metabolite bioproduction as a function of pH. pH of the MTC-5 medium (5 g/L cellobiose) was adjusted to indicated value and log-phase culture of each strain was used to inoculate each pH treatment. Sampling was performed at 24 h. Control refers to unadjusted pH of the medium (pH 6.9). At pH 6, none of the strains were able to grow.
